# Ecological character displacement destabilizes food webs

**DOI:** 10.1101/824615

**Authors:** Matthew A. Barbour

**Affiliations:** University of Zurich, Department of Evolutionary Biology and Environmental Studies, Winterthurerstrasse 190, 8057 Zurich, Switzerland; University of British Columbia, Department of Zoology, 2212 Main Mall, Vancouver, BC V6T 1Z4, Canada

**Keywords:** competition, eco-evolutionary dynamics, consumer-resource interactions, adaptation, community stability

## Abstract

Ecological character displacement is an adaptive process that generally increases phenotypic diversity. Despite the fact that this diversification is due to an eco-evolutionary feedback between consumers competing for shared resources, its consequences for food-web dynamics have received little attention. Here, I study a model of two consumers competing for two shared resources to examine how character displacement in consumer attack rates affects resource abundances and the resilience of food webs to perturbations. I found that character displacement always strengthened consumer-resource interactions whenever consumers competed for resources that occurred in different habitats. This increase in interaction strength resulted in lower resource abundances and less resilient food webs. This occurred under different evolutionary tradeoffs and in both simple and more realistic foraging scenarios. Taken together, my results show that the adaptive process of character displacement may come with the ecological cost of decreasing food-web resilience.

## Introduction

Ecological character displacement is an important adaptive process in generating biodiversity (Schluter, 2000; Pfennig & Pfennig, 2010). This process is due to “phenotypic evolution in a species generated or maintained by [exploitative] resource competition with one or more coexisting species” (Schluter, 2000). A large body of theoretical (e.g. Lawlor & Smith, 1976; Abrams, 1986; Doebeli, 1996; Taper & Chase, 1985; McPeek, 2019) and empirical (reviewed in: Schluter, 2000; Dayan & Simberloff, 2005; Stuart & Losos, 2013) work has examined which scenarios lead to phenotypic divergence or convergence of competing consumers. The general conclusion has been that, if resources are nutritionally substitutable (Abrams, 1987; Fox & Vasseur, 2008) and there is no other strong source of density dependence acting on consumers (Abrams, 1986), then resource competition drives the adaptive divergence of competitors (Lawlor & Smith, 1976; Taper & Chase, 1985). This adaptive process is not simply a response to static differences in resource distributions, but creates an eco-evolutionary feedback that drives further differentiation. This crucial insight was made by theoretical models that explicitly included resource dynamics as a mediator of competition in driving evolutionary change (Lawlor & Smith, 1976; Abrams, 1986; Taper & Chase, 1985).

Although models that included resources led to insights about the evolution of character displacement, the ecological feedback onto consumer-resource dynamics has received surprisingly little attention. This is likely because the ecological feedback has been primarily studied through the lens of coexistence theory (Lawlor & Smith, 1976; Germain *et al.*, 2018; Bassar *et al.*, 2017; McPeek, 2019). For example, early theoretical work showed that character displacement promotes coexistence by favoring specialized consumers that experience reduced interspecific competition (Lawlor & Smith, 1976). Yet, this reduction in interspecific competition may, at the same time, increase interspecific interactions between specialized consumers and their resources. Both food-web theory and empirical studies have shown that increasing the strength of consumer-resource interactions often suppresses the abundance of resources, which if sufficient enough, can generate oscillations and less stable consumer-resource dynamics (Rosenzweig, 1971; Luckinbill, 1973; Murdoch *et al.*, 2002, 2003; McCann, 2011). Thus, a food-web perspective, which accounts for both the direct and indirect effects of consumer-resource interactions, may yield new insight to the ecological consequences of character displacement.

Here, I address this knowledge gap by studying a mathematical model that examines how ecological character displacement affects consumer-resource dynamics in a food-web context. Specifically, I sought to answer the question: how does character displacement in consumer attack rates affect resource abundances and food-web stability? To test the generality of these effects, I explored different ecological foraging scenarios and evolutionary tradeoffs in consumer attack rates. I found that the adaptive process of character displacement often comes with an ecological cost; resulting in food webs with lower resource availability and that are less resilient to perturbations.

## Material and methods

### Underlying consumer-resource dynamics

To examine how ecological character displacement affects resource abundances and food-web stability, I analyzed a continuous-time model of two consumers (*C*_*j*=1,2_) competing for two shared resources (*R*_*i*=1,2_):

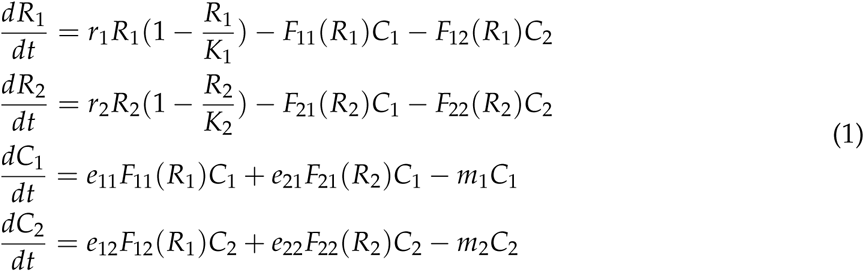

where *r*_*i*_ represents the intrinsic growth rate of resource *i, K*_*i*_ represents the carrying capacity of resource *i, e*_*ij*_ represents the conversion efficiency of resource *i* into consumer *j*, and *m*_*j*_ represents the mortality rate of consumer *j. F*_*ij*_(*R*_*i*_) represents consumer *j*’s feeding rate on resource *i* (i.e., its functional response). This model is a useful characterization of a scenario where consumers compete for two distinct resources (e.g. zooplankton and benthic invertebrates in lakes) rather than a scenario where resources are better characterized by a continuous trait distribution (e.g., seed size; see Taper & Chase (1985) for an example). Importantly, inferences about character displacement can only be made by comparing food webs with and without a competing consumer (Schluter & McPhail, 1992). Therefore, I arbitrarily set *C*_2_ = 0 to create a food-web without a competing consumer for these comparisons.

### Foraging scenarios

I studied three different foraging scenarios. In the first, I assume that consumers can forage for both resources simultaneously (Fig. 1a) and their feeding rate increases linearly with resource abundance, such that:

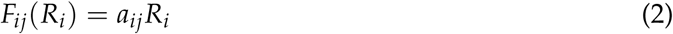

where *a*_*ij*_ is the attack rate of consumer *j* on resource *i*. This first scenario is the starting point for many models of resource competition (MacArthur, 1972); however, it does not reflect many food webs where consumers are mobile and their foraging behavior links resources that occur in different habitats (McCann *et al.*, 2005). The second scenario accounts for this spatial context (Fig. 1b) and takes the form:

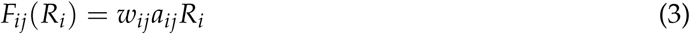

where *w*_*ij*_ represents the proportion of time consumer *j* spends foraging in a habitat where only resource *i* is found (i.e., its habitat preference). Note that since *w*_*ij*_ is a proportion that *w*_1,*j*_ = 1 − *w*_2,*j*_. Finally, it is well known that consumer feeding rates often saturate at high resource abundances (Holling, 1959; Rosenzweig & MacArthur, 1963; Murdoch *et al.*, 2003; McCann, 2011) and that consumers do not usually spend a fixed proportion of time in a particular habitat (McCann *et al.*, 2005). The third scenario accounts for these biological realities and takes the form (derived by McCann *et al.*, 2005):

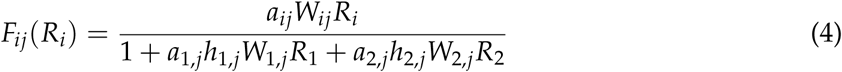

where consumer *j*’s feeding rate on resource *i* is influenced by the abundance of each resource; saturates as resource abundances increase (due to handling time *h*_*ij*_); and consumer habitat preferences are modified by the relative abundance of resources, such that: 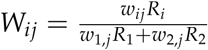.

**Figure 1:**
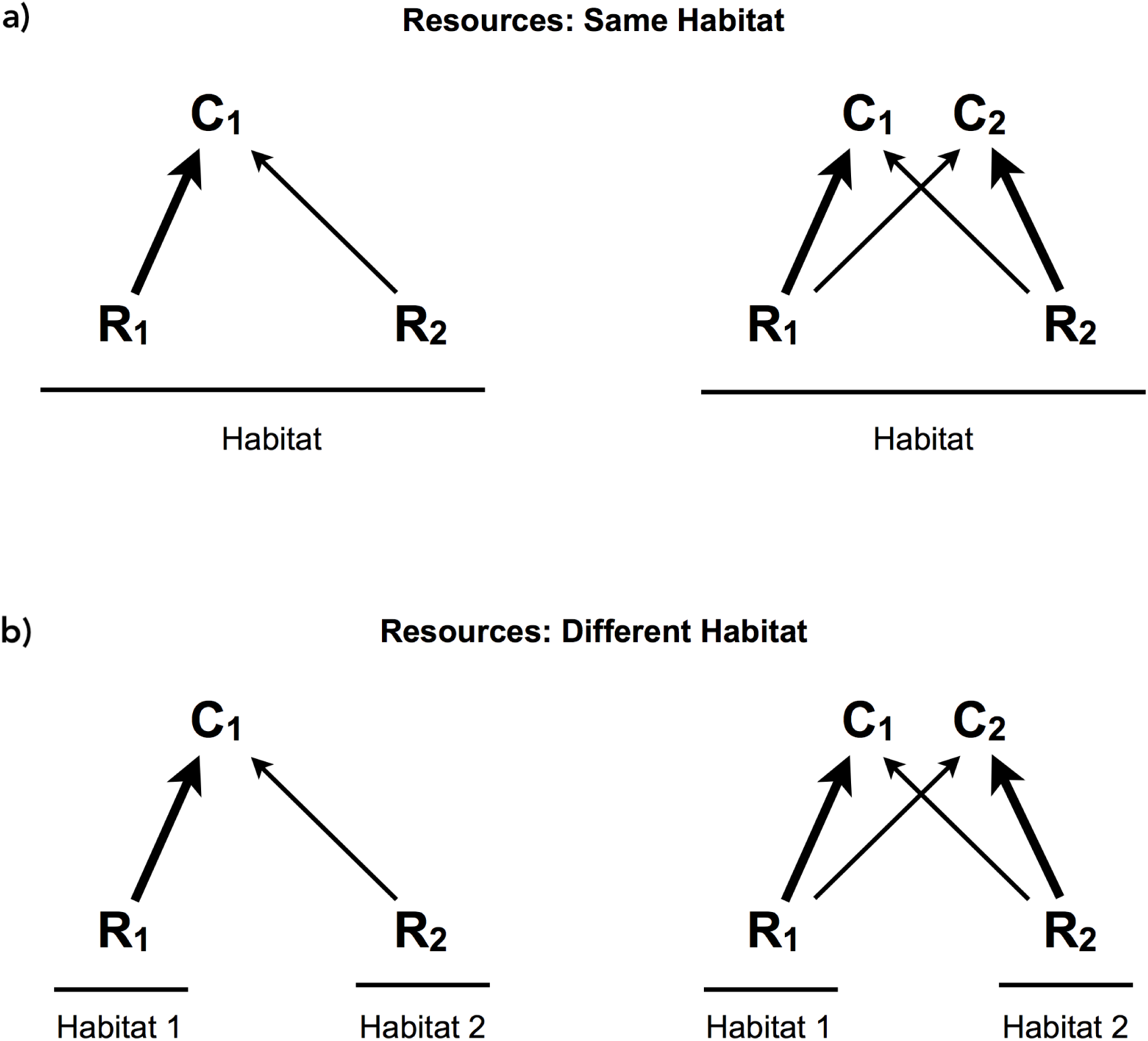
Ecological foraging scenarios. I examined whether the effect of ecological character displacement on food-web dynamics depended on whether consumers competed for resources that occurred in the same (a) vs. different habitats (b). Note that inferences about character displacement can only be made by comparing food webs with (right) and without (left) a competing consumer, so I arbitrarily set *C*_2_ = 0 for these comparisons. The width of each arrow corresponds to the initial attack rate (*a*_ij_) of consumer *j* on resource *i*. Note that *C*_1_ was pre-adapted to *R*_1_ (*a*_11_ > *a*_21_), while *C*_2_ was a mirror image, being pre-adapted to *R*_*2*_ (*a*_22_ = *a*_11_; *a*_12_ = *a*_21_). In each scenario, I assumed consumer feeding rates increased linearly with resource abundance. I also relax this assumption and consider a more realistic functional response when resources occurred in different habitats (b).

Previous studies have analyzed the evolution of consumer attack rates in the first two foraging scenarios using an Adaptive Dynamics approach, with the general result being divergent character displacement (Lawlor & Smith, 1976; Abrams, 1986). I also used an Adaptive Dynamics approach to analyze the evolution of consumer attack rates in the third foraging scenario, and I too observed divergent character displacement (detailed analysis given in Appendix S1). I say consumers have undergone divergent character displacement if their evolved attack rates are more specialized when evolving with vs. without a competing consumer. Specialization of consumer *j* on resource 1 is measured as 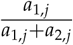, where a value of 0.5 is a complete generalist (*a*_1,*j*_ = *a*_2,*j*_), and a value of 1 is a complete specialist (*a*_2,*j*_ = 0). Values less than 0.5 indicate specialization on the other resource. Since I did not observe convergent character displacement in any of the foraging scenarios I analyzed, I refer to divergent character displacement as simply (ecological) character displacement throughout the rest of the text.

### Food-web dynamics

Given that character displacement occurred across these foraging scenarios, I focus here on its consequences for food-web dynamics. To do this, I analyzed differences in resource abundances and food-web stability at equilibrium. An equilibrium is reached when there is no change in the population growth rates of consumers and resources (i.e., the rates of change in equation 1 are 0), and solving the system at this point gives equilibrium abundances for each resource 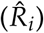 and each consumer (*Ĉ*_*j*_). I also compared the local stability of these food webs using standard methods (Otto & Day, 2007). This stability analysis derives the dominant eigenvalue, *λ*_max_, of the matrix of partial derivatives of each species’ population growth rate (given by equation 1) with respect to each species’ abundance evaluated at equilibrium. If −*λ*_*max*_ > 0, then the food web will return to equilibrium after a small perturbation (i.e., it is locally stable), with more positive values indicating a faster return time. If −*λ*_*max*_ < 0, then the food web is not locally stable.

When possible, I derived analytical expressions for the relationship between consumer attack rates and food-web dynamics. To do this, I simplified the model by assuming that resources are equivalent (*r* = *r*_*i*_ and *K* = *K*_*i*_) as well as consumers (*e* = *e*_*ij*_; *h* = *h*_*ij*_; *m* = *m*_*j*_), except that consumer attack rates and their habitat preferences (if present) are mirror images of each other (*a*_11_ = *a*_22_; *a*_12_ = *a*_21_; *w*_11_ = *w*_22_). Note that I arbitrarily set *C*_1_ as being pre-adapted to *R*_1_ (*a*_11_ > *a*_21_; *w*_11_ > 0.5), and therefore *C*_2_ was pre-adapted to *R*_2_. Controlling for other sources of variability allowed me to isolate the general effects of character displacement. All mathematical derivations were conducted in Mathematica (Wolfram Research Inc., 2018) and are provided in Appendices S1-3.

To gain insight to the eco-evolutionary feedback generated by character displacement, I conducted simulations using an Adaptive Dynamics approach. Specifically, after letting consumer and resource abundances reach a steady state, I created a mutant consumer by randomly choosing one and modifying its attack rate on one resource by either subtracting or adding a small constant (0.01 in the following simulations) with equal probability. The mutant’s attack rate on the other resource was determined by a tradeoff, such that (*a*_1,*j*_/*A*)^*n*^ + (*a*_2,*j*_/*A* ^*n*^)= 1, where *A* is the total investment in attack rates and *n* describes the shape of the tradeoff (Sargent & Otto, 2006). This function has the useful property that it differentiates between cases where intermediate combinations of *a*_1,*j*_ and *a*_2,*j*_ are higher than the extremes (when *n* > 1, green line in Fig. 2) or, conversely, where the two extremes are higher than intermediate investments (when *n* < 1, orange line in Fig. 2). When *n* = 1, the tradeoff function is linear, and all combinations of *a*_1,*j*_ and *a*_2,*j*_ have the same total attack rate (blue line in Fig. 2). Assuming the mutant consumer was rare, I then determined whether the mutant had higher relative fitness than the resident consumer, and thus could invade and replace the resident consumer. If the mutant was able to invade, I updated the attack rate of the resident consumer to the mutant attack rate and allowed consumer and resource abundances to reach a steady state. I then repeated the simulation up to 10,000 times, which was sufficient for consumers to either reach an evolutionary stable strategy (ESS, Smith & Price, 1973) or an evolutionary limit (e.g., 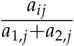 is constrained to a maximum of 1 and minimum of 0). Unless otherwise noted, I conducted simulations with the following parameter values: *r* = 1; *K* = 4; *e* = 0.8; *m* = 1; *A* = 2; *h* = 0.4; and *w*_11_ = *w*_22_ = 0.6. I set an initial value of *a*_11_ = *a*_22_ = 1.2, while *a*_12_ and *a*_21_ depended on the value of *n*. I set initial consumer and resource abundances to: *R*_1_ = *R*_2_ = 2; *C*_1_ = *C*_2_ = 1. All simulations were conducted in R (R Core Team, 2018) and the code to reproduce these simulations is publically available on GitHub (https://github.com/mabarbour/ECD_model) and has been archived with Zenodo (https://zenodo.org/badge/latestdoi/79349192).

**Figure 2:**
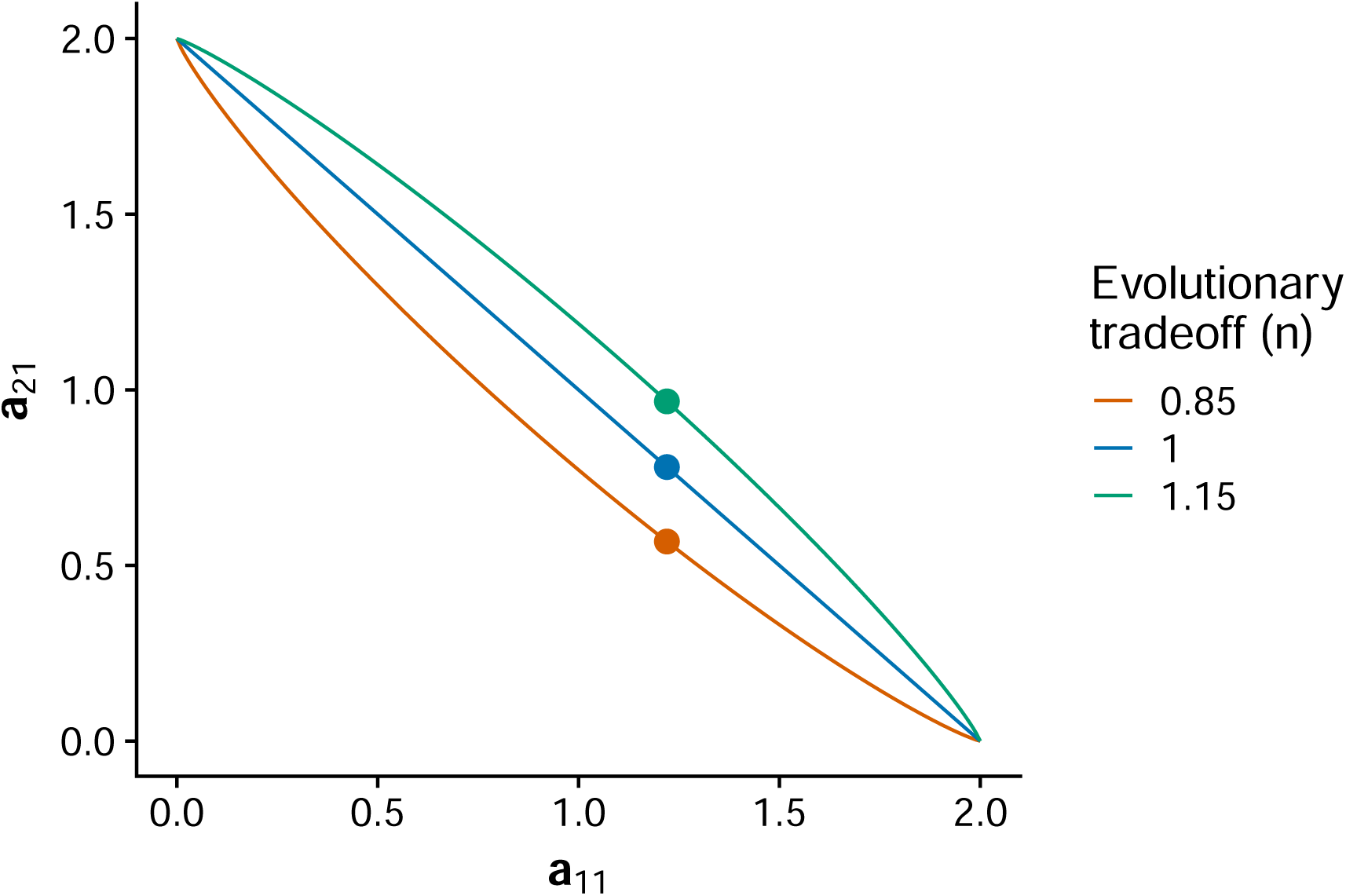
Evolutionary tradeoffs in consumer attack rates. In each foraging scenario, I explored the effects of three different tradeoffs: intermediate combinations of attack rates (*a*_1,*j*_, *a*_2,*j*_) are higher than the extremes (green line, *n* > 1); extreme combinations of attack rates are higher than intermediate investments (orange line, *n* < 1); and all combinations of attack rates have the same total attack rate (blue line, *n* = 1). Points corresponding to attack rates at the beginning of the simulation for *C*_1_, which was pre-adapted to *R*_1_ (*a*_11_ > *a*_12_). Note that *C*_2_ was a mirror image of *C*_1_, being pre-adapted to *R*_2_ (*a*_22_ = *a*_11_; *a*_12_ = *a*_21_).

## Results

### Resources occur in the same habitat

In this first scenario (equation 2), the abundance of resources at equilibrium are equivalent when both consumers and resources are present 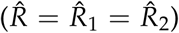, and are determined by the following equation (derived in Appendix S2):

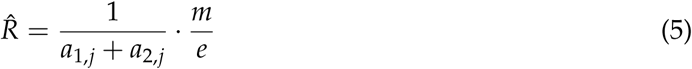

A key determinant of resource abundance in this scenario is the consumer’s total attack rate, *a*_1,*j*_ + *a*_2,*j*_. Therefore, the effect of character displacement on food-web dynamics depends on how the shape of the tradeoff function influences the evolution of consumer attack rates.

I found that the shape of the tradeoff function qualitatively affects the relationship between character displacement and resource abundances in this scenario (Fig. 3a,b). For example, if consumer’s are constrained by a linear tradeoff (blue lines), then there is no net change in total attack rate (Fig. 3a) and character displacement has no effect on resource abundances (Fig. 3b). If the tradeoff is concave down (green lines), then resource abundances can actually increase under character displacement (Fig. 3b). This is because the total attack rate of consumers is maximized at intermediate values (*a*_1,*j*_ = *a*_2,*j*_) and decreases as consumers diverge (Fig. 3a). When the tradeoff is concave up (orange lines), character displacement suppresses resource abundances due to the increase in total attack rates (Fig. 3a,b). Although the equation I derived for resource abundances was for the scenario where both consumers and both resources were present, it accurately predicts the abundance of resources when a single consumer reaches its evolutionary stable strategy (ESS; triangles on respective colored lines in Fig. 3b). This is because a single consumer evolves to be a generalist that has equal attack rates on each resource (triangles at 0.5 along x-axis in Fig. 3a), resulting in equivalent resource abundances.

**Figure 3:**
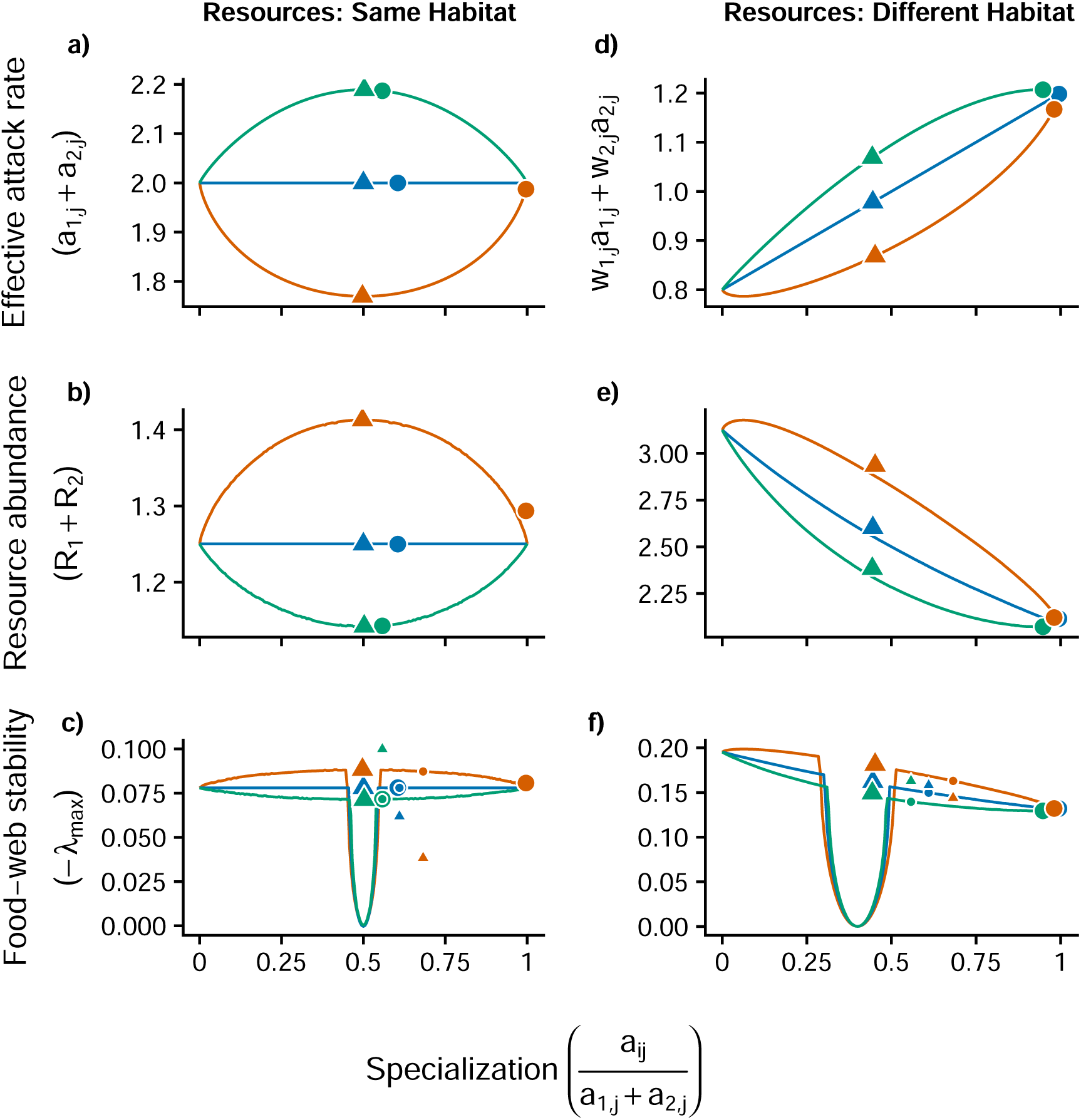
Effect of character displacement on food-web dynamics under different evolutionary tradeoffs and foraging scenarios. Lines show predicted values when both consumers and resources are present. Different line colors correspond to different tradeoffs in attack rates (green, *n* = 1.15; blue, *n* = 1; orange, *n* = 0.85). Large circles (two consumers) and triangles (one consumer) correspond to the end points of the eco-evolutionary simulation for *C*_1_ (the choice to display *C*_1_ was arbitrary), whereas as small shapes correspond to the starting points (only in stability panels). In both foraging scenarios, feeding rates increase linearly with resource abundance, but the equation for the effective attack rate is different.

The effect of character displacement on resources corresponds to its impact on food-web stability. For example, when character displacement decreases resource abundances (orange points in Fig. 3b), there is also a decrease in food-web stability (Fig. 3c). Character displacement may not affect or even increase food-web stability (blue and green lines in Fig. 3c); however, evolution does not favor strong divergence in these scenarios (blue and green points in Fig. 3a), which dampens these contingent effects. Note that the dip in stability occurs when both consumers evolve to be generalists, a situation that is not favored in any of the foraging scenarios I examined (Fig. 3c).

### Resources occur in different habitats

In the second foraging scenario (equation 3), I again see that resource abundances are equivalent when both consumers and resources are present 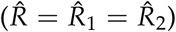, but are now determined by the following equation (derived in Appendix S3):

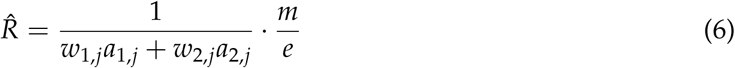

This equation implies that if consumers evolve to become specialists on resources that occur in their preferred habitat (e.g., *w*_1,*j*_ > 0.5 and *a*_1,*j*_ > *a*_2,*j*_), then the effective attack rate of consumers (*w*_1,*j*_*a*_1,*j*_ + *w*_2,*j*_*a*_2,*j*_) will always increase, regardless of the tradeoff (Fig. 3d). Thus, character displacement always results in resource suppression (Fig. 3e). Note that the shape of the tradeoff can modify the effect of character displacement. This is not so much due to the tradeoff affecting the magnitude of displacement (it does, but the effect is minor), but because the form of the tradeoff affects resource abundances when a single consumer has reached an ESS (triangles in Fig. 3e). In contrast, resource abundances reach a similar value when consumers evolve in the presence of a competitor (circles in Fig. 3e), because character displacement tends to reach a constraint of complete specialization. It is worth noting that resource abundances are consistently higher at the single consumer ESS compared to the predictions I derived for when both consumers are present (deviation of triangles from respective colored lines in Fig. 3e). This is because consumers actually evolve slightly specialized attack rates on the resources that occur in their non-preferred habitat (deviation of triangles from 0.5 along x-axis in Fig. 3d).

As seen previously, the effect of character displacement on resource abundances qualitatively corresponds to its effect on food-web stability (Fig. 3f). Specifically, character displacement decreases food-web stability, regardless of the tradeoff in attack rates. This is not simply a consequence of having an additional consumer in the system, but emerges from the eco-evolutionary feedback between character displacement and resource suppression (Fig. 3c). For example, when the tradeoff is concave up (orange), the initial two-consumer food web (small circle) is more stable than when there is only one consumer (small triangle); however, this pattern switches by the end of the eco-evolutionary simulation (large points).

### Adding a more realistic functional response

In the third foraging scenario (equation 4), I observed the same general effect of character displacement as the previous scenario (resources in different habitats, but linear functional response). This is because resource abundances at equilibrium are governed by a similar dynamic (derived in Appendix S1):

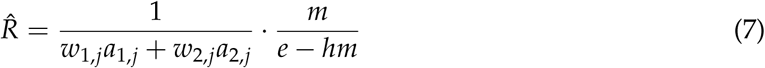

And since evolution favors character displacement toward their preferred resources (see Appendix S1), the effective attack rate of consumers (*w*_1,*j*_*a*_1,*j*_ + *w*_2,*j*_*a*_2,*j*_) will always increase, resulting in lower resource abundances and decreased food-web stability (Appendix S4, Fig. S1).

In the first two foraging scenarios, character displacement influences food-web stability, but all of the food webs ultimately return to a stable equilibrium (because −*λ*_*max*_ > 0, see Appendix S2-3). In this more realistic model, however, whether the food web is locally stable depends on consumer and resource parameters. Specifically, I found that the two-consumer food web will transition from having a locally stable equilibrium to a limit cycle under the following conditions (derived using Routh-Hurwitz criteria in Appendix S1):

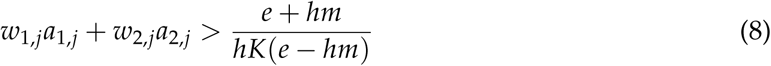

This inequality indicates that character displacement always pushes the food web toward an unstable structure in this more realistic foraging scenario (Fig. 4). Note that I stopped the simulation in the two-consumer food web once it became locally unstable. I do not simulate beyond this point as this would require making assumptions about the dynamics of mutant consumers in variable environments, which is beyond the scope of this work.

**Figure 4:**
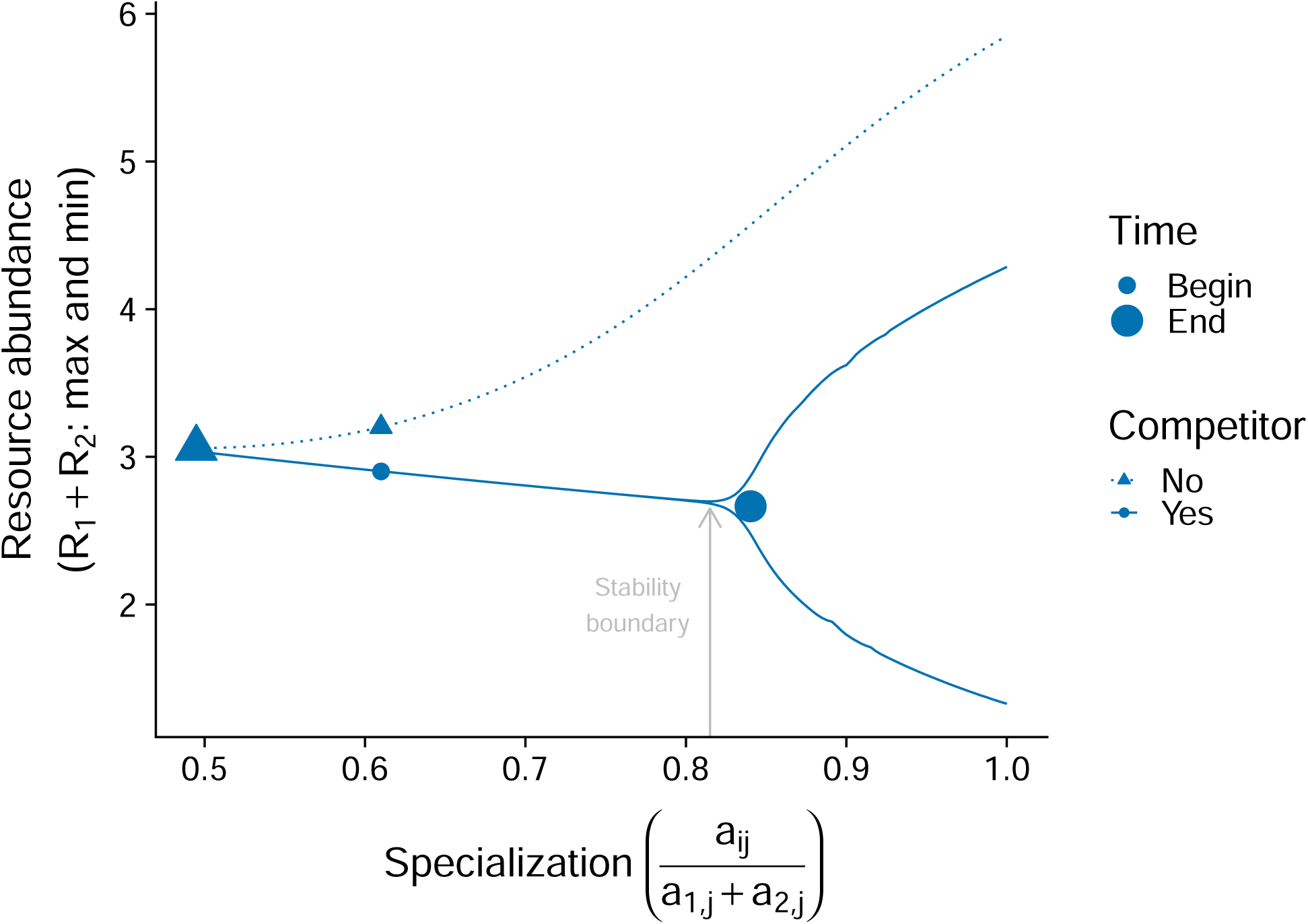
Character displacement creates an unstable food web. Lines illustrate the effect of character displacement across the range of specialization for *C*_1_ (the choice to display *C*_1_ was arbitrary), while the points are the results of an eco-evolutionary simulation. Note that I increased the total investment in attack rates (*A* = 3.3) to create a scenario that could result in an unstable food web. Although I specified a linear tradeoff in attack rates for this simulation, different tradeoff shapes do not qualitatively alter these results (see Appendix S4, Fig. S1).

### Robustness to consumer asymmetry

The previous analytical results and simulations make a strong assumption that competing consumers start off as perfect mirror images of each other (i.e., there is symmetry). Yet, theory indicates a predictable asymmetry between initial consumer attack rates. This predictable asymmetry emerges from a process of community assembly where a single consumer invades a system, evolves to be a generalist that can equally attack both resources, followed by the invasion of a second, more specialized, consumer. This theoretical scenario has been hypothesized as the sequence of events leading to character displacement in threespine stickleback in small coastal lakes of British Columbia (Schluter & McPhail, 1992; Schluter, 2000).

To test whether my results were robust to this asymmetry, I used the evolved attack rates at the end of the simulations with one consumer as the starting values for one of the two consumers. I did this for all foraging scenarios and tradeoffs previously examined. I found that my previous inferences are robust to including consumer asymmetry across different foraging scenarios and tradeoffs (Appendix S4, Fig. S2-3).

## Discussion

### Resource abundances

One of the criteria used to demonstrate ecological character displacement is that “sites of sympatry [two consumers] and allopatry [one consumer] should not differ greatly in food [resource abundances]” (Schluter & McPhail, 1992). In contrast, my results indicate that character displacement causes predictable differences in resource abundances. In fact, the ecological and evolutionary scenarios that favored the largest character displacement always decreased the relative abundance of resources. For example, if mobile consumers compete for resources that occur in different habitats, then character displacement always resulted in lower resource abundances. Threespine stickleback, one of the classic examples of character displacement, exemplify this foraging scenario (Schluter & McPhail, 1992; Schluter, 2000). Stickleback must move between the pelagic and littoral zones of a lake when foraging for zooplankton and benthic invertebrates, respectively. The theory developed here predicts that resource abundances will be lower in lakes where competing stickleback have undergone character displacement compared to lakes with only a single species of stickleback. Interestingly, a disproportionate number of the documented cases of character displacement involve carnivores (Schluter, 2000) that are larger, and likely more mobile, than their resources (McCann *et al.*, 2005), suggesting that many cases of ecological character displacement may result in lower resource availability.

Similarly, the evolutionary tradeoff that favored character displacement decreased resource availability across all foraging scenarios. Although data on the shape of the tradeoff in consumer foraging traits is scarce, two classic examples of character displacement, Darwin’s finches and threespine stickleback, both appear to exhibit a tradeoff where extreme trait values increase the net foraging rate of consumers (Schluter *et al.*, 1985; Arnegard *et al.*, 2014). While it is theoretically possible that character displacement does not alter (or even increase) resource abundances, this was limited to the simplest, and arguably least realistic, foraging scenario and under tradeoffs that did not favor large displacements, and thus less likely to detect in nature. Taken together, my results call for empirical work to test these clear theoretical predictions and suggest a revision is needed for one of the criteria used to demonstrate character displacement.

### Food-web stability

My most striking result was that ecological character displacement made food webs less resilient to perturbations. In fact, under the most realistic foraging scenario, character displacement can even result in an unstable food web. The mechanism underlying this destabilization is quite general. Character displacement generally increases the strength of consumer-resource interactions, but does not alter the strength of intraspecific interactions. This relative increase in interspecific interactions, combined with the natural oscillatory tendency of consumer-resource dynamics (Lotka, 1925; Volterra, 1926), creates a food-web structure that is less resilient to perturbations (Chesson & Kuang, 2008; Rip & McCann, 2011; McCann, 2011).

Interestingly, the ecological conditions that favor character displacement are those that are already the least resilient to perturbations. For example, McPeek (2019) showed that character displacement is favored in food webs that are either highly productive, easy to find and capture resources, or under weak abiotic stress. This corresponds to higher values of *K* (productivity) or *A* (investment in attack rates), or lower values of *m* (abiotic stress). Each of these corresponding changes decrease food-web resilience, as they increase the strength of consumer-resource interactions relative to intraspecific interactions. For example, increasing productivity reduces intraspecific competition in resource populations while increasing the flux of energy to consumers, resulting in the paradox of enrichment (Rosenzweig, 1971). Similarly, higher feeding rates or lower consumer mortality both increase the relative strength of consumer-resource interactions, which predictably destabilizes food webs (Rip & McCann, 2011; McCann, 2011). This suggests that the most dramatic examples of character displacement will not only occur in, but also cause, the least stable food-web structures.

A handful of empirical patterns support the hypothesis that character displacement decreases food-web resilience. For example, a single species of threespine stickleback lives in hundreds of small coastal lakes in British Columbia, but the species pair, where character displacement has resulted in specialized limnetic and benthic species, are only known from six lakes in four independent watersheds (Schluter & McPhail, 1992; Schluter, 2000). Perhaps many lakes had a species pair in the past, but have lost a species due to a less resilient food-web structure (Borrelli, 2015; Borrelli *et al.*, 2015). The species pair are known to be vulnerable to perturbations, as they have gone extinct in two of the six lakes after the introduction of nonnative species (Hatfield, 2001; Taylor *et al.*, 2006; Rudman & Schluter, 2016). The vulnerability of the stickleback system also corresponds with the fact that aquatic food webs have several properties that make them less resilient to perturbations, such as higher productivity and more efficient energy transfer to consumers (Rip & McCann, 2011). Detecting the ghost of competition past (Connell, 1980) may be quite difficult, but it could be possible with recent advances in genomics. For example, Feulner & Seehausen (2019) detected genomic signatures of hybridization in sympatric whitefish species following periods of eutrophication. Perhaps solitary stickleback in some lakes retain genomic signatures of having been a habitat specialist in the past.

My results contrast, but do not necessarily contradict, the notion from coexistence theory that character displacement contributes to species coexistence (Lawlor & Smith, 1976). Rather than studying resilience, coexistence theory usually studies the mutual ability of consumers with different phenotypes to invade when rare (i.e., mutual invasibility; Chesson, 2000). In the context of character displacement, a shortcoming of this mutual invasibility measure is that it does not allow a comparison between food webs with and without a competing consumer. Such comparisons are necessary for inferring the effects of character displacement, a point that has been made clear in the criteria to demonstrate character displacement (Schluter & McPhail, 1992; Schluter, 2000). Although the addition of a consumer to a food web can decrease its resilience in the absence of evolution (May, 1973), my results are primarily driven by an eco-evolutionary feedback between consumer evolution and resource abundances.

### Caveats

Although I model the indirect effects of coevolution between consumers, I do not account for potential coevolution between consumers and resources. In the context of my model, I would expect prey to evolve traits that reduce consumer attack rates. Thus, prey evolution would act to counter the effects of character displacement on resource abundance and food-web stability. Note that this does not negate my general conclusion that ecological character displacement decreases resource abundances and stability; however, this process may itself create another eco-evolutionary feedback between consumers and resources. This may actually help maintain dramatic examples of character displacement and prevent them from destabilizing systems, because it allows consumer traits to become decoupled from their attack rate. Examining this decoupling would be ideal in a quantitative genetic model that explicitly tracks trait dynamics, but it would not fundamentally change the conclusions presented here.

Another potential caveat is that I explored my model in a setting that makes many assumptions about resource and consumer symmetry (but see consumer asymmetry section). Prior work has shown that allowing for resource asymmetry, for example, may decrease the magnitude of character displacement (Abrams, 1986). While this may dampen the amount of divergence, it should not qualitatively change the relationship I observed.

I studied this eco-evolutionary feedback between consumers and resources using an Adaptive Dynamics approach. A strength of this approach is that it enabled me to gain analytical insight to the effects of character displacement in a more realistic foraging scenario. This is much less tractable in quantitative genetic (Taper & Chase, 1985; McPeek, 2017) or explicit genetic (Doebeli, 1996) models of character displacement, which is why the foraging scenarios previously examined have been limited (but see McPeek, 2017). A weakness, however, is that I assume a separation of time scales between ecological and evolutionary dynamics, an assumption that is becoming less tenable (Hairston *et al.*, 2005; Hendry, 2016). I also do not explicitly model an underlying phenotypic trait for consumer attack rates nor do I allow for intraspecific variation. That being said, my theoretical predictions are likely robust to these assumptions. This is because models that explicitly include resource dynamics inevitably show that resource competition results in character displacement, regardless of whether a quantitative genetic or Adaptive Dynamics approach is used (Lawlor & Smith, 1976; Taper & Chase, 1985). A quantitative genetic model may certainly show differences in the pace of character displacement, but this should not qualitatively change its effect on food-web dynamics. It is important to note that my conclusions only apply to food webs with biotic resources that are nutritionally substitutable. It would be interesting to extend these current analyses to non-substitutable resources where convergent character displacement is expected (Abrams, 1987; Fox & Vasseur, 2008).

### Conclusions

Here, I show that an adaptive process that generates phenotypic diversity generally makes that diversity more susceptible to future extinctions. This destabilizing effect emerges from an eco-evolutionary feedback involving direct and indirect interactions between species in a food-web context. This result contrasts with the current notion that patterns of phenotypic diversity are solely the result of evolutionary constraints imposed by mutation, natural selection, gene flow, and genetic drift. In particular, my result supports the recent suggestion that food-web stability can impose an ecological constraint on phenotypic diversity that is agnostic to these evolutionary processes (Borrelli *et al.*, 2015). I expect that identifying when and where this ecological constraint arises will yield novel insight to the patterns of biodiversity we see in nature.

## Acknowledgements

This work was inspired by discussions with Seth Rudman, Dolph Schluter, Ben Gilbert, and Kevin McCann. Sally Otto provided much help in early analyses of the mathematical model. I would not have been able to take the theory as far as I did without her guidance and encouragement. Matt Osmond, Jean Gibert, and Seth Rudman provided critical feedback on an earlier draft. For funding support, I thank the University of British Columbia (Four-Year Fellowship to M.A. Barbour), NSERC (Discovery grant to Greg Crutsinger), and the Swiss National Science Foundation (grant 31003A_160671 to Jordi Bascompte).

