## Appendix S4 for "Ecological character displacement destabilizes food webs"

### Ecological character displacement destabilizes food webs

Matthew A. Barbour[1,2,*]

1. University of Zurich, Department of Evolutionary Biology and Environmental Studies, Winterthurerstrasse 190, 8057 Zurich, Switzerland;

2. University of British Columbia, Department of Zoology, 2212 Main Mall, Vancouver, BC V6T 1Z4, Canada;

*Elements*: Figures S1, S2, and S3.

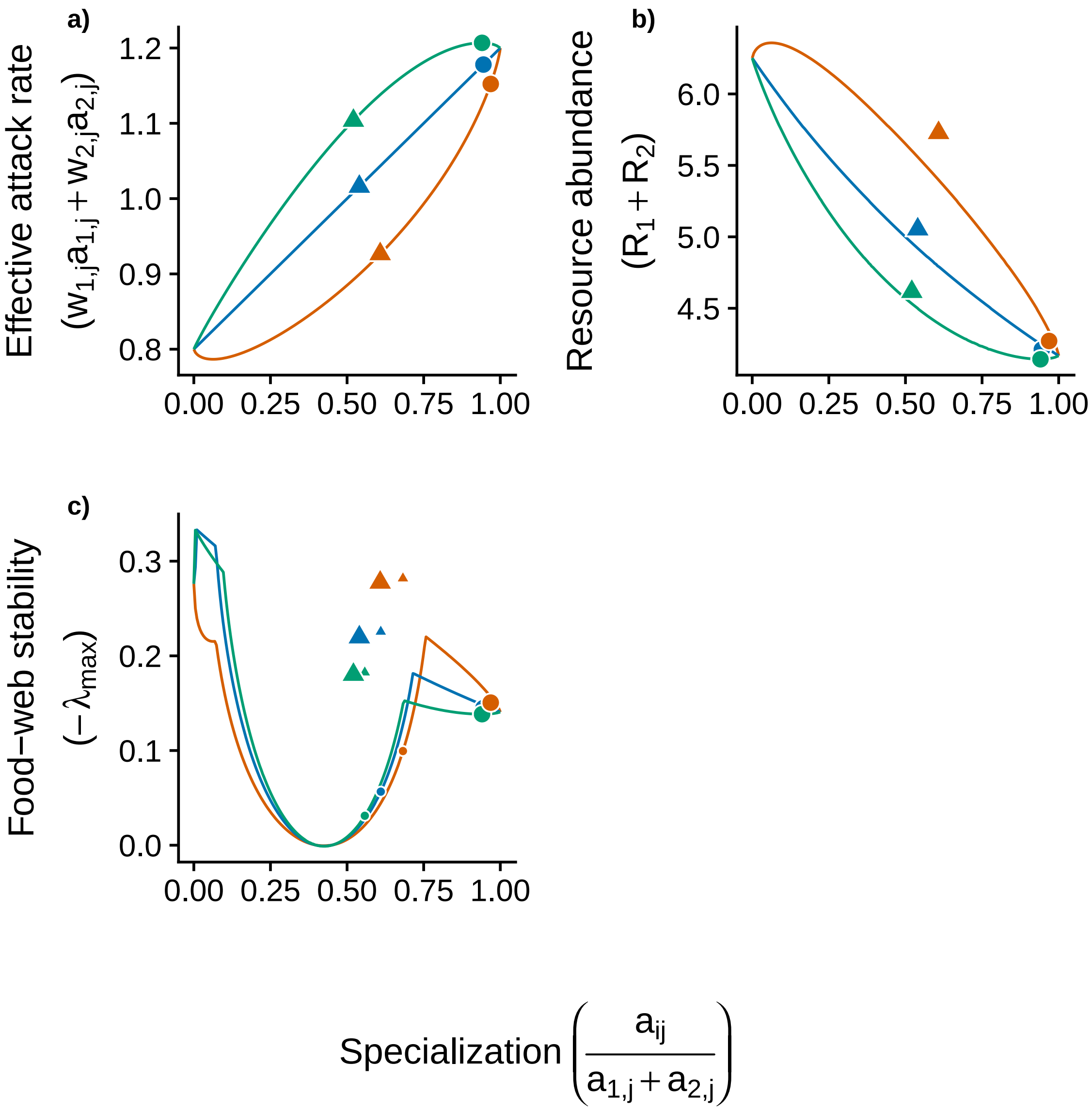


Figure S1: **Effect of character displacement when consumers exhibit a more realistic functional response.** Different line colors correspond to different tradeoffs in attack rates (green, $n$ = 1.15; blue, $n$ = 1; orange, $n$ = 0.85). Large circles (two consumers) and triangles (one consumer) correspond to the end points of the eco-evolutionary simulation for $C_1$ (choice to display $C_1$ was arbitrary), whereas as small shapes correspond to the starting points (only in stability panel). This figure illustrates how, regardless of the tradeoff, character displacement increases the effective attack rate of consumers. This always resulted in a suppression of resource abundances and concomitant decrease in food-web stability. Initial parameter and state variables are given in the main text.

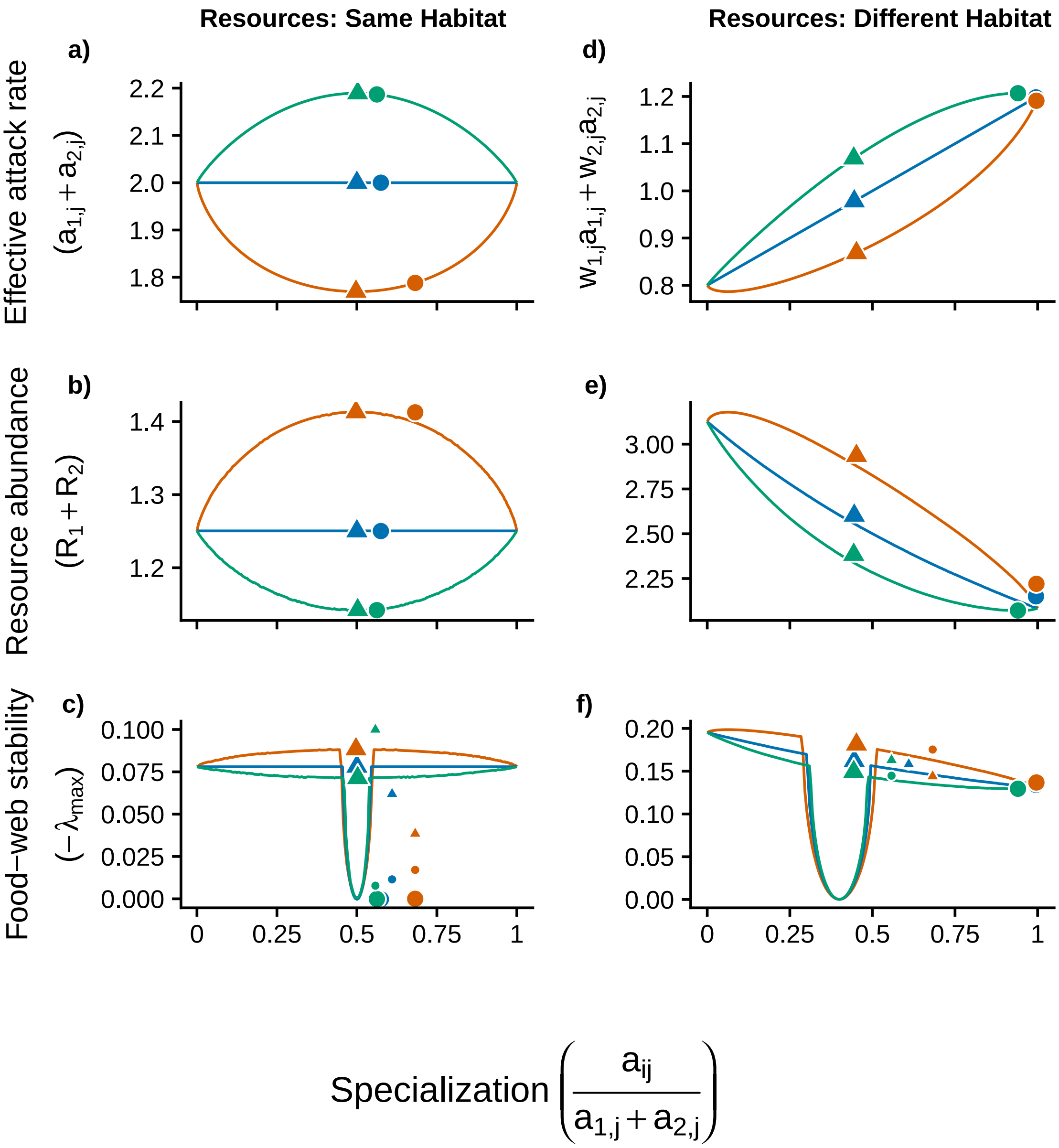


Figure S2: **Robustness to consumer asymmetry**. Lines show predicted values when both consumers and resources are present. Different line colors correspond to different tradeoffs in attack rates (green, $n = 1.15$; blue, $n = 1$; orange, $n = 0.85$). Large circles (two consumers) and triangles (one consumer) correspond to the end points of the eco-evolutionary simulation for $C_1$ (choice to display $C_1$ was arbitrary), whereas as small shapes correspond to the starting points (only in stability panels). In both foraging scenarios, feeding rates increase linearly with resource abundance, but the equation for the effective attack rate is different. The similarity with Fig. 3 (main text) shows that adding an asymmetry in consumer attack rates does not alter the conclusions reported in the main text. If anything, it is more likely that an asymmetry will push the system toward the boundary of stability (note position of large circles in lower left stability panel). Details of the consumer asymmetry simulation are given in the main text.

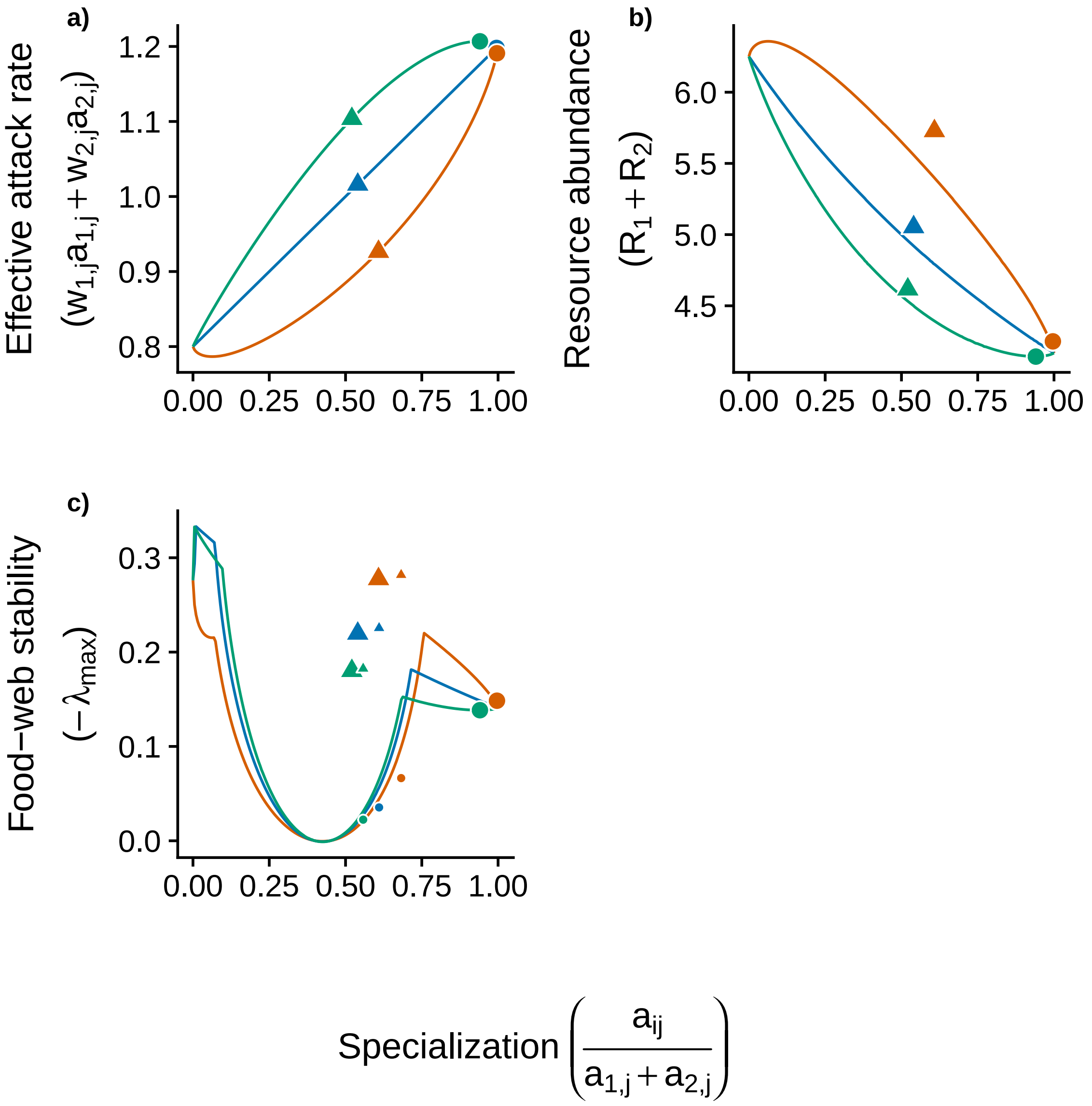


Figure S3: **Robustness to consumer asymmetry when consumers exhibit a more realistic functional response.** Different line colors correspond to different tradeoffs in attack rates (green, $n$ = 1.15; blue, $n$ = 1; orange, $n$ = 0.85). Large circles (two consumers) and triangles (one consumer) correspond to the end points of the eco-evolutionary simulation for $C_1$ (choice to display $C_1$ was arbitrary), whereas as small shapes correspond to the starting points. Regardless of the tradeoff, character displacement increases the effective attack rate of consumers (a). This always resulted in a suppression of resource abundances (b) and concomitant decrease in food-web stability (c). The similarity with Fig. S1 shows that adding an asymmetry in consumer attack rates does not alter the conclusions reported in the main text. Details of the consumer asymmetry simulation are given in the main text.
